# RNA polymerase II depletion from the inactive X chromosome territory is not mediated by physical compartmentalization

**DOI:** 10.1101/2021.03.26.437188

**Authors:** Samuel Collombet, Isabell Rall, Claire Dugast-Darzacq, Alec Heckert, Aliaksandr Halavatyi, Agnes Le Saux, Gina Dailey, Xavier Darzacq, Edith Heard

## Abstract

Sub-nuclear compartmentalization has been proposed to play an important role in gene regulation by segregating active and inactive parts of the genome in distinct physical and biochemical environments, where transcription and epigenetic factors are either concentrated or depleted. The inactive X chromosome offers a paradigm for studying sub-nuclear compartmentalization. When the non-coding *Xist* RNA coats the X chromosome, it recruits repressors and chromatin factors that trigger gene silencing, and forms a dense body of heterochromatin from which the transcription machinery appears to be excluded. Phase separation has been proposed to be involved in X-chromosome inactivation (XCI) and might explain exclusion of the transcription machinery by preventing its diffusion into the *Xist*-coated territory. Here, using quantitative fluorescence microscopy and single particle tracking, we show that RNA polymerase II (RNAPII) freely accesses the *Xist* territory during initiation of XCI, and that its diffusion is not prevented by biophysical constraints. Instead, the apparent depletion of RNAPII is due to the loss of its chromatin bound fraction. These findings demonstrate that initial exclusion of RNA Pol2 from the inactive X is a consequence of its reduced binding rate at the chromatin and gene level, rather than the biophysical compartmentalization of the inactive X heterochromatin domain. The Xist silent compartment is thus a biochemical rather than a biophysical compartment, at least during initiation of XCI.

## Introduction

In female eutherian mammals, one of the two X chromosomes becomes silenced through the process of XCI. This is controlled by the non-coding *Xist* RNA which becomes up-regulated from one of the two X chromosomes during early development, coating the chromosome *in cis* and triggering chromosome-wide gene silencing. One of the earliest events occurring during XCI is the very rapid exclusion of the transcriptional machinery from the *Xist* coated part of the X-chromosome territory, both in early embryos *in vivo* (*1*) and in early differentiating mouse embryonic stem cells (mESCs) (*2*). This has led to the suggestions that the Xi may form a specific sub-nuclear compartment which could constitute a biophysical “barrier” preventing RNAPII and other transcription factors from entering the *Xist*-coated territory, in order to ensure and/or to maintain gene silencing (*3*). Indeed X-linked genes become relocated into the Xist compartment as they become silenced, whilst genes that escape XCI remain located outside it (*2*) (*4*), although whether this gene relocation is a cause or a consequence of X-chromosome gene silencing events is still not clear.

*Xist* is a multi-tasking molecule that recruits many proteins via its conserved repeats to the future inactive X chromosome (Xi). This includes SPEN, which is brought via the *Xist* A repeats and is essential for gene silencing (*5*),(*6*). The PRC1 Polycomb complex is also recruited via *Xist* B repeats, while PRC2 gets accumulated on the Xi downstream of this (*7*) (*8*) (*9*) (*10*). Other factors are also recruited via *Xist* E repeats, including CIZ1 at the onset of XCI (*11*), and PTBP1, MATR3, and CELF1 at later stages (*12*). These proteins become highly accumulated on the inactive X in a *Xist* dependent manner. Protein-RNA and protein-protein interactions have been proposed to allow their accumulations through liquid-liquid phase separation, forming so-called membraneless organelles such as the nucleolus or stress granules. A number of *Xist* recruited factors, including SPEN, PRC1 and PTBP1-MATR3, form low affinity protein-protein interactions (*12, 13*), suggested to increase molecular crowding in the *Xist* compartment and participate in the compaction of chromatin (*13*). Accordingly, phase separation has been proposed to underlie the compartmentalization of the inactive X (*14*). This hypothesis would offer a simple mechanism for the exclusion of RNAPII, as *Xist* would trigger the formation of a heterochromatic membraneless organelle from which the transcription machinery is physically excluded. However, other mechanisms have been shown to allow apparent nuclear compartmentalization, such as modulation in protein-DNA binding rate, without involving a phase separation process (*15*). To investigate the nature of the *Xist* RNA nuclear compartment and its relationship to RNAPII, we have applied quantitative and super-resolution fluorescence microscopy to assess the concentrations and the dynamics of RNAPII in living cells, both inside and outside the *Xist* compartment as it forms and gene silencing initiates.

### Results

#### RNAPII concentration within the *Xist* RNA compartment

In order to assess the dynamics and nature of the RNAPII-depleted *Xist* RNA compartment, we took advantage of a mouse embryonic stem cell line (mESC) in which the endogenous *Xist* gene on one chromosome is controlled by a doxycycline inducible promoter (*16*). This allows for rapid and synchronous induction of *Xist* expression and X-chromosome inactivation (XCI). To visualize *Xist* RNA and RNAPII in live cells, we tagged *Xist* (on the doxycycline inducible allele) with BGL stem loops that can be specifically bound by a Bgl-G protein fused to a GFP (*17*) (*6*), and in the same cells, we generated homozygous HaloTag knock-ins (KI) at the endogenous RNA Polr2a and Polr2c genes, encoding the RPB1 and RPB3 proteins respectively, which are two major subunits of RNAPII (**Fig 1A** and **Fig S1A-B**). Both the RPB1- and RPB3-Halo KI cell lines displayed slightly higher levels of the tagged proteins compared to wild-type cells (**Fig S1E**) probably due to a stabilisation of the protein by the tag. After 24h *Xist* induction, which was previously shown to be sufficient for global silencing of almost all X-linked genes (*18*) (*6*), these cells show efficient *Xist* RNA coating (in >50% of cells) and X-linked gene silencing (**Fig S1F-G**). Using confocal microscopy, we image *Xist*-Bgl-GFP and RNAPII-Halo in 3D (with 0.48 microns between stacks, see methods) in live cells following 24h of *Xist* induction, and systematically segment the *Xist* compartment (XC), the nucleoplasm and nucleoli (as a control regions where RNAPII is excluded) using robust machine learning segmentation tools (*19*) (**Fig 1B and S2A-C**).

**Fig 1.**
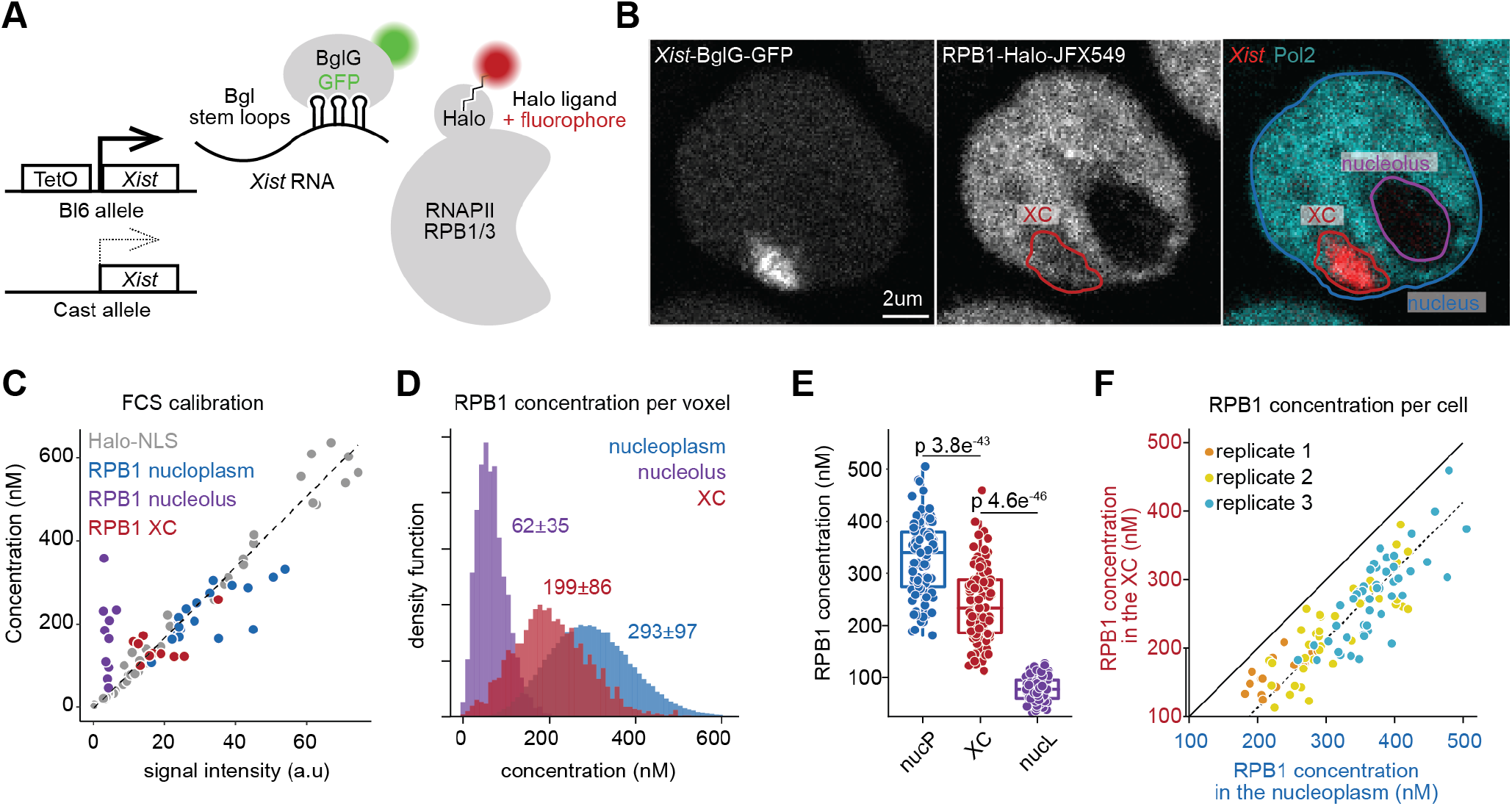
RNAPII concentration in the *Xist* compartment. **A.** scheme of Xist and RNAPII tagging for combined live cell imaging. **B.** Representative image of confocal microscopy of Xist (BglG-GFP) and RNAPII (RPB1-Halo) in live cells (single Z stack) after 24h of Xist induction (doxycycline treatment) with overlaid segmentation of nucleus, nucleoli and Xi (see methods). **C.** Calibration of signal intensity from point scanning imaging with FCS measured concentrations. Each dot represents a single measurement from a single cell. The linear calibration is established only on the freely diffusing Halo-NLS. Note that fluorescence intensity in the nucleolus was below the threshold of robust FCS measurement (see supplementary figure S2D), leading to artifactual concentration estimation. **D.** Calibrated RNAPII (RPB1) concentration per voxel for the nucleus shown in B, based on the calibration in C. The average concentration per region is indicated (± 95% confidence interval). **E.** distribution of average concentration per region per cell after 24h of Xist induction. Each dot represents a single cell (n=107). P-values of the differences are indicated on top (t-test two sided, paired data). **F** RPB1 Concentration in the XC versus nucleoplasm. Each dot represents a single cell.

We then assessed whether RNAPII was completely excluded from the *Xist* RNA territory region or whether it could still be detected within the *Xist* domain. We used fluorescence correlation spectroscopy (FCS) calibrated imaging (*20*), where FCS measurements on a freely diffusing control (Halo-NLS) allow calibrating fluorescence intensity from confocal images into absolute concentration (**Fig 1C and Fig S2E**). FCS measurements for RNAPII either inside the *Xist* RNA territory or in the nucleoplasm, followed the same linear trend as the freely diffusing Halo-NLS control (**Fig 1C and Fig S2D**), indicating that RNAPII concentration in these regions could be robustly measured. We then used this calibration curve to convert 3D image voxel intensity to local concentration (**Fig 1D and Fig S2F**) and found that the concentrations of RPB1 were significantly lower in the XC compared to the nucleoplasm (**Fig 1E**), with an average reduction of-88 nM. RPB1 levels nevertheless remain substantial within the *Xist* territory, with an average concentration of 207nM (corresponding to an average 1,165 molecules of RPB1 per XC). Similar results were found for RPB3 with an average concentration of 220nM (average of 934 molecules perXC) (**Fig S2G**). Interestingly, the concentration of RPB1 and RPB3 in the *Xist* compartment scaled linearly with their nucleoplasmic concentration (**Fig 1F and S2H**) showing a constant and consistent offset deficit, implying that the reduction in RNAPII concentration on the XC is not mediated by a sequestration away from this chromosome territory. As virtually all X-linked genes are silenced after 24h *Xist* induction, these results suggest that gene silencing on the inactive X is not mediated by a physical exclusion of RNAPII from the *Xist* territory.

#### RNAPII can diffuse freely through the *Xist* compartment

We next more directly addressed whether the reduction in RNAPII concentration on the XC was due to a ‘barrier effect’ reducing RNAPII flux exchanging in and out of the domain. We performed single particle tracking (SPT) using a photo-activable Halo-PA-JF646, and high speed imaging (5.477ms frame interval and 1ms excitation pulse) to capture the trajectories of diffusing molecules (**Fig 2A**) (*21, 22*). We then compared the flux of RNAPII molecules entering the Xi territory to comparable regions in the nucleoplasm (see methods) (**Fig 2B**). Because XC territories have different sizes and shapes we first compared the distribution of entering events and found no significant difference for both RPB1 and RPB3 (Kolmogorov-Smirnov test pValue=0.45 and 0.56, respectively) (**Fig 2C**). At the single cell level, RNAPII flux is strongly correlated to that in the rest of the nucleoplasm (**Fig 2D**), The differences in concentration observed earlier (**Fig 1G**) can therefore not be attributed to a potential domain exclusion mechanism such as phase separation (Cerase et al. 2019). Finally, we wondered whether Pol2 molecules entering the XiC are more likely to return to the nucleoplasm (**Fig 2E**). Looking at the distribution of angles of entering trajectories, we did not observe a clear difference inside and outside XiC, for both RPB1 nor RPB3 (**Fig 2F**). Quantifying the ratio of forward (0 to 30 degrees) and backward angles (160 to 180 degrees) as previously done (*22*), (*23*), we saw no difference between the XiC and shifted regions for RPB1 (**Fig 2G**) and only a slight reduction for RPB3 (forward/backward=0.92), suggesting that the trajectories of Pol2 molecules entering the XiC are not significantly affected.

**Fig 2.**
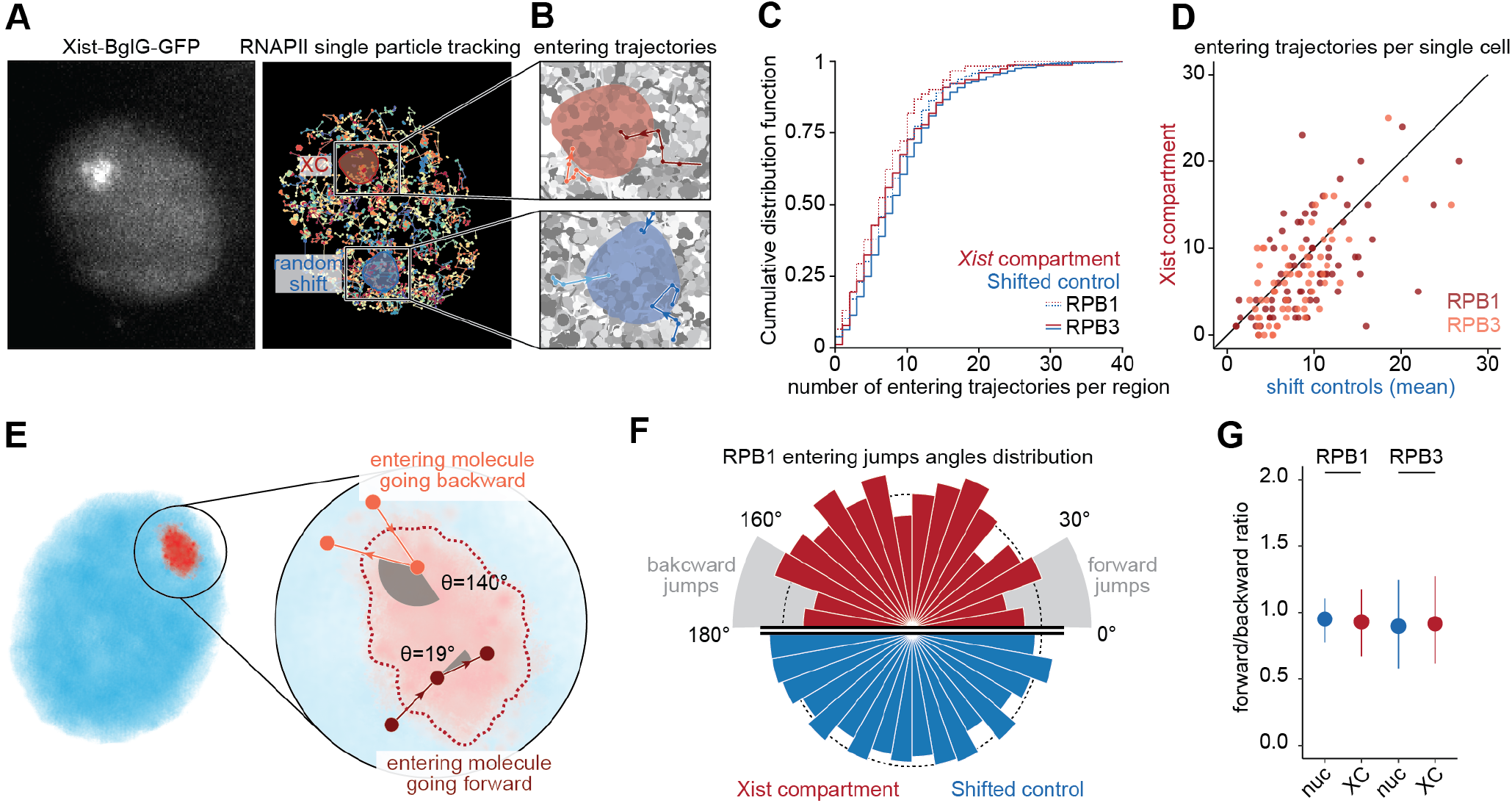
Characterisation of RNAPII flux with XC. **A.** Example of Xist-BGL-GFP imaging and RNAPII RPB1 single particle tracking (5.477ms between frame, 1ms exposure, 3 minutes tracking) in the same cell. Each trajectory is shown with random colors. The red area shows the segmentation of the Xist compartment, and the blue area a random spatial shift of this region (taking into account the distance to the nuclear periphery, see methods). **B.** Zoom into the XC region and random shift to highlight entering trajectories of RNAPII into the XC. **C.** Cumulative distribution function of the number of entering trajectories in the XC (red) and random shifts (blue) for RPB1 (plain line) and RPB3 (dashed line). Only trajectories with a mean square root displacement (MSRD) >200nm were selected to ensure to use only freely diffusing molecules (see methods). **D.** Scatter plot of the number of trajectories entering the XC (y-axis) and the average number of trajectories entering the shift controls (x-axis) in the same single cell, for RPB1 (red) and RPB3 (orange). **E.** If a molecule “bounces back”, its trajectory should display large angles between jumps, while molecules that “move forward” should display small angles. Molecules in Brownian motion in free space should show no preference. **F.** Distributions of entering trajectories angles between the entering jump and the following one (as depicted in **E**) for RPB1 trajectories entering the XC (red, n=556 entering jumps) or shifted control regions (blue, n=4,408 entering jumps). The radius of the bar represents the density of counts. **G.** Ratio of forward angles (0 to 30 degrees) / backward angles (160 to 180 degrees) as previously done (22, 23). The dot represents the fraction estimated from all pooled trajectories (n=556/4,408 and 346/3310 trajectories entering XC/shift control region, for RBP1 and RPB3 respectively), and the error bar represents the standard deviation from 50 bootstrap subsampling of 250 entering jumps (see methods).

Altogether, these results show that RNAPII can freely enter the *Xist* compartment.

#### RNAPII apparent exclusion is due to the loss of its stably bound fraction

Given these results, we hypothesized that the apparent decrease in RNAPII concentration (**Fig 1E**) could actually be the result of a loss of the bound fraction of RNAPII on the *Xist* RNA coated inactive X chromosome, while the diffusing fraction is at an equilibrium with the nucleoplasm. Fitting a two component model (bound vs freely diffusing) to our SPT data (**Fig 3A**), we found a significant reduction in the bound fraction on the XC for both RPB1 and RPB3 (**Fig 3B**). Scaling the concentration measured by FCS-CI (**Fig 1F**) by the bound/free fraction measured by SPT (**Fig 3C**), we confirmed that the amount of freely diffusing RNAPII is not significantly different between the XC and the nucleoplasm (in agreement with no impairment in RNAPII diffusion) but the amount of bound RNAPII is reduced. Yet approximately 1/3 of RNAPII molecules on XC are bound presumably specifically immobilized on chromatin (**Fig S4A-B**). Because fast SPT is not ideal to interrogate long binding events as observed with elongating RNAPII in the nucleoplasm (**Fig S4C**) (*21*) we investigated further this bound fraction using FRAP. Using FRAP we found that almost all fluorescence on the XC was recovered one minute after photobleaching (**Fig 3D**) while only 63% was recovered on a region of the same size in the nucleoplasm, showing that the 33% “bound” fraction on XC revealed by SPT corresponds to RNAPII molecules only transiently immobile, and that the “stably” bound fraction (approximately 40% of RNAPII in the nucleoplasm) is completely lost. This result clearly shows that RNAPII binding on the Xi is so transient that it cannot account for any significant elongation and therefore is probably restricted to non-promoter interactions and/or abortive promoter interactions. Scaling the FRAP signal in the XC to the nucleoplasmic level showed that similar absolute levels are recovered in both regions, corresponding to the initial levels of RNAPII on the XC. This indicates that virtually all RNAPII on the XC correspond to the same amount of freely diffusing and transiently bound RNAPII as in the nucleoplasm. Altogether, these results imply that the reduction of RNAPII on the *Xist* compartment is due to the loss of initiation and a partial loss of pre-initiation.

**Fig 3.**
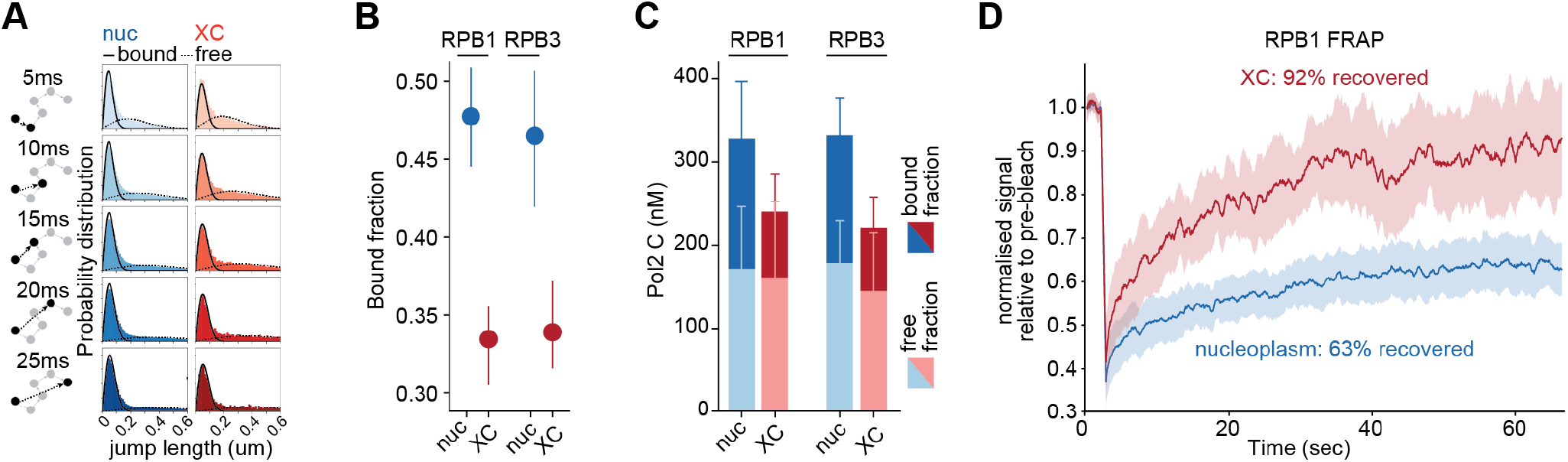
RNAPII dynamics on XC. **A.** Fitting of a two component model (bound versus freely diffusing molecules) to the distribution of jump length at different time scales, for trajectories inside the XC or outside (nuc), pooling all data from 96 single cells. **B.** Estimates of the bound fraction inside and outside the XC, for RPB1 and RPB3. The dot represents the fraction estimated from all pooled trajectories, and the error bar the standard deviation from 50 bootstrap subsampling of 3,000 trajectories (see methods). **C.** Concentration of RPB1 and RPB3 (from Fig 1F and S2G) scaled by the bound and free fraction from B. The error bar represents the 95% confidence interval for the product of concentration and bound/free fraction. Calculated using the delta method (24) (see methods). **D.** Fluorescence recovery after photobleaching (FRAP) experiment for the XC or control nucleoplasmic region. Signal is normalized to the signal before bleaching (see methods). The line represents the mean of signal (n=15 cells) and the shade its 95% confidence interval.

#### RNAPII diffusion is not altered in the *Xist* territory

Finally, we investigated whether the diffusion of RNAPII was affected in a qualitative manner on a *Xist* territory. The inactive X has previously been shown to be more compact than its active counterpart (~ 1.2 fold compaction) (*25, 26*). In addition, *Xist*-recruited proteins such as SPEN, Ciz1 and PRC1 have recently been shown to form supra-molecular complexes upon their recruitment on the inactive X (*13*). Increased molecular crowding, due to protein complexes and/or higher chromatin density, could constrain the diffusion of RNAPII (**Fig 4A**). Looking at the jump angle distribution for free RNAPII (MSRD > 200nm), we did not observe any difference for RPB1 trajectories inside XC or in shifted control region (**Fig 4B and C**), and only a slightly lower forward/backward ratio for RPB3 (0.83 in XC versus 0.97 in shifted control region, **Fig 4C and S5a**). This result suggests that molecular crowding on the *Xist* territory has only a minor effect (if any) on RNAPII diffusion. Accordingly, we did not measure a significant difference in diffusion coefficient inside and outside the XC (**Fig 4D and S5b**).

**Fig 4.**
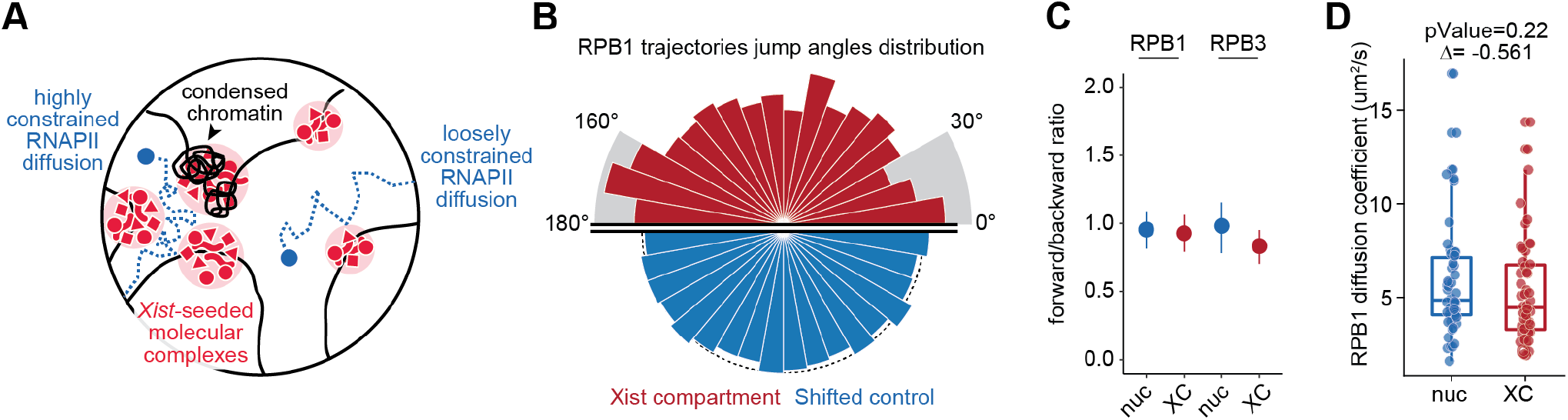
RNAPII diffusion on the Xi. **A.** Schematic representing how different environments might constrain RNAPII (in blue) diffusion. On the left side, Xist-seeded molecular complexes (in red) and dense chromatin (black) occupy a significant space which constrain RNAPII diffusion. On the right side, protein complexes and chromatin do not occupy a significantly higher space and RNAPII diffusion is not impacted. **D.** Distributions of jump angles for RPB1 (tops) and RPB3 (bottom) trajectories entering the XC (red) or control regions (blue). The radius of the bar represents the density of counts. **C.** Ratio of forward angles (0 to 30 degrees) / backward angles (150 to 180 degrees). The dot represents the fraction estimated from all free trajectories (MSRD>200nm, n=1410/13,335 and 834/8,826 trajectories in XC/shift, for RBP1 and RPB3 respectively), and the error bar represents the standard deviation from 50 bootstrap subsampling of 500 free trajectories (see methods). **D.** RPB1 diffusion coefficient based on FCS measurements inside and outside the XC. Each dot represents a single measurement in a single cell (measurement inside and outside are paired per single cell).

Together, these results show that neither chromatin compaction nor *Xist* RNA associated molecular complexes affect RNAPII diffusion during XCI initiation.

### Discussion

The sequestering functions of molecular compartmentalisation in the nucleus have been long-proposed to play a role in the regulation of gene expression. Local modifications in the concentrations of regulatory proteins, through biochemical or physical compartmentalization, may affect the efficiency of the processes they regulate. While this seems to be the case for repressors recruited by *Xist*, such as SPEN, PTBP1 and MATR3 (*12*) (Markaki et al. 2020), we show here that RNAPII diffusion is not prevented at the level of the all *Xist*-coated territory, resulting in a similar concentration of free RNAPII as in the nucleoplasm (**Fig 5A**). The inactive X chromosome has previously been shown to be a heterogeneous structure (*27*), formed of distinct, tightly packed heterochromatin domains (*28*). In addition, *Xist*-associated proteins have been shown to form supra-molecular complexes through protein-protein interactions, which was proposed to increase molecular crowding and participate in chromatin compaction (Markaki et al. 2020). Indeed, molecular condensates have been the subject of much attention recently.Phase separated protein domains are seen as regions where multivalent protein-protein interactions can modulate mass action, tweaking the “on rates” of transcription factors binding to DNA (*29*). Here we show that the nature of the multivalent assemblies decorating Xi chromatin during initiation of XCI does not affect the target search process or concentration (on rate) of free RNAPII molecules (**Fig 5B**). Rather, they act simply by preventing RNAPII binding to its DNA binding sites on chromatin, resulting in a loss of the stable bound fraction on the *Xist* territory (**Fig 5C**). Taken together, our results suggest that the exclusion of RNAPII from the inactive X chromosome is not caused by a homogenous biophysical compartment-sequestering function, but rather by a modification of its chromatin that protects/represses RNAPII binding events. This is in agreement with the fact that the multivalent protein complexes involved in X inactivation decorate small chromatin domains in a manner not compatible with the formation of a large phase separated whole chromosome domains (*13*).

**Fig 5.**
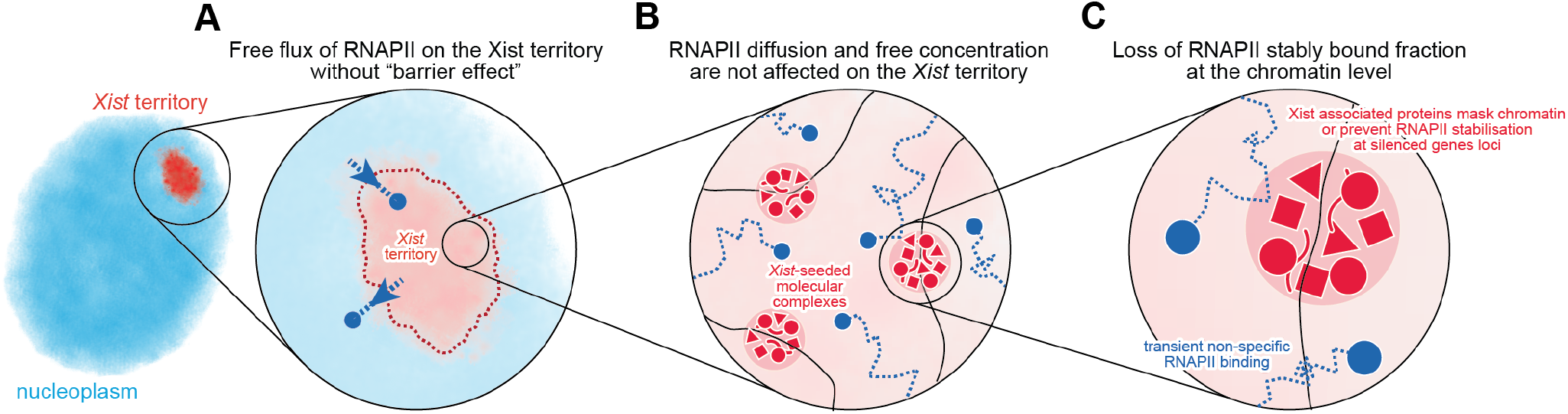
Summary of RNAPII dynamics on the *Xist* territory at multiple scales.

Two mechanisms can be proposed to explain how *Xist*-associated factors prevent stable binding of the RNAPII: i) they mask its binding sites on chromatin and impede RNAPII access, or ii) they prevent RNAPII stabilization at its target sites. Our FRAP data show that the amount of transiently bound RNAPII is the same inside and outside the XC (**Fig S4F**), therefore suggesting that *Xist* and its associated factors do not significantly prevent RNAPII access to DNA, but only its stabilization. In addition, the acute depletion of SPEN ((*6, 30*) or the deletion of *Xist* A-repeats (that are required for SPEN recruitment) ((*6, 30*)) result in a near complete loss of gene silencing, but do not affect *Xist* global accumulation on the X chromosome. However, deletion of the *Xist* A-repeats prevents its coating of transcriptionally active regions of the genome (*31*), suggesting that chromosomal *Xist* coating and the formation of supra-molecular complexes are are not necessary for the initiation of gene silencing. Together, these results suggest that *Xist* RNA and its associated proteins do not act by physically masking genes, but rather by inhibiting RNAPII transcription at the pre-initiation or initiation stage. Accordingly, inhibition of the assembly of the pre-initiation complex via triptolide results in the loss of RNAPII stable binding, but not its transient binding (*23, 32*), similar to what we observe on the *Xist* territory. Overall, these results suggest that the mechanism of gene silencing on the inactive X does not involve a mechanism of biophysical compartmentalization, but rather relies on the same fundamental principles mediating gene regulation on autosomes, i.e. the biochemical modulation of transcription initiation.

The depletion of RNAPII within the *Xist* territory, which was described as one of the first events during XCI (*1*)(*2*), is still observed upon induction of a *Xist* mutant lacking its A-repeats and therefore unable to silence genes (*2*). This can be explained by the fact that in the absence of the A-repeat, as *Xist* cannot spread fully over active regions of the X chromosome (*31*),these remain at the periphery of or outside the *Xist* territory (*2*). In this case the *Xist* compartment consists of only silent regions of the chromosome, including most repetitive sequences (*4*). Here, RNAPII is not expected to bind stably, and this therefore results in a reduction in concentration within the *Xist* domain, as visualized by RNAPII immunofluorescence or live imaging, or by Cot-1 RNA FISH.

The conclusions we have drawn here, focus only on the initial steps of XCI when Xist coats the X chromosome and gene silencing begins. Once *Xist* has triggered XCI, other factors and chromatin modifications accumulate on the inactive X including, in some cases quite late during differentiation, such as the PTBP1-MATR3 complex, at around day 4 (*12*). Furthermore, the inactive X chromosome has been shown to occupy a smaller volume (~1.2 fold) than its active counterpart in differentiated cells in culture and in vivo blastocyst stage embryos (*33*) (*25*) (*26*). Indeed, recent reports suggest that this compaction occurs only after several days of differentiation (*13*), probably through the late recruitment of the SMCHD1 protein (*34*) (*35*). These factors could affect RNAPII diffusion and lead to the formation of a biophysical compartment. Therefore, while our results show that impeded RNAPII diffusion within the *Xist* coated domain of the X is not required for the initiation of gene silencing, it could still play a later role in the maintenance of XCI.

## Acknowledgements

The authors would like to thank Luke Lavis for providing all the Halo-ligand dye used in this study; all members of the Advanced Light Microscopy Facility and the Cytometry Facility at EMBL for their help; Robert Tjian, Francois Dossin, Marko Lampe, Arina Rybina and Merle Hantsche-Grininger for their help and their feedback on the manuscript. The work performed in E.H. lab was supported by an ERC Advanced Investigator Award ERC-ADG-2014 671027. The work performed in X.D. lab was financed by the NIH 1U54CA231641 grant. S.C. is supported by an EMBO long-term fellowship (EMBO ALTF 275-2018) and supported by the Joachim Herz Foundation.

## Material and Methods

### Cell culture and treatment

Mouse XX ES cells (TX1072) were grown on 0.1% gelatin-coated flasks in 8% CO2 at 37C and cultured in mouse ES media serum+2i+LIF: DMEM (Sigma) without phenol-red, 15% FBS (Gibco), L-Glutamin (584 mg/L), non-essential amino-acids (ThermoFisher #15140122, 6mL/L), sodium pyruvate (110mg/L), 0.1 mM β-mercaptoethanol, 1,000 U ml−1 leukaemia inhibitory factor (LIF, Merck ESG110), CHIR99021 (3 μM), PD0325901 (1 μM. *Xist* expression was induced upon administration of doxycycline (2 μg/ml).

### CRISPR-Cas9 genome editing

#### Transfection

All transgenic insertions were performed using the 4D nucleofector system from Lonza. For each nucleofection, five million cells were electroporated with in presence of the plasmids (MidiPreps). For targeted knock-in (RBP1- and RPB3-Halo, BglG-GFP at Tigre), 2.5ug of non-linearized targeting vectors and 2.5ug of sgRNA were used. For Halo-NLS and H2b-Halo, 2.5ug of piggybac construct containing the transgene and 2.5ug of transposase plasmid were used.

#### Selection

For BglG-GFP knock-in at Tigre, the insert also contained a puromycin selection cassette. After nucleofection cells were splitted into 10cm with serial dilution (1/10, 1/100 and 1/1000). 48h after cells seeding puromycin was added to culture media (0.4 μg/mL). After 1 week of selection, single clonal colonies were picked (see *Clonal expansion* here after).

#### FACS sorting

after transfection cells were then put back in culture in a T25 for 2 days, passage once in a T75 and culture 2 more days. Cells were labelled using Halo-ligand-JF646 (graciously provided by Luke Lavis) at 100nM in media, incubated for 30min, washed 3 times in PBS, incubated 3 times in fresh media for 20min with PBS washes between incubations. Cells were dissociated using Accutase(Invitrogen), washed twice in medium, and resuspended in sorting buffer. 1000 positive cells were then sorted on a FACSAria fusion. Cells were then put back in culture in a 10cm petri dish coated with gelatin for 7 to 10 days before picking clones.

#### Clonal expansion

Single clonal colonies were manually picked under an EVOS cell imaging system, incubated in trypsin for 10min, and splitted 3/4 - 1/4 in two 96-well plates coated with gelatin and cultured for 2 days. The high density plate was used for PCR genotyping and the low density plate for clone expansion.

### DNA and RNA Pyro-sequencing

DNA was extracted using DNeasy Blood and Tissue kit. RNA extraction was performed using the RNeasy kit and on-column DNase digestion (Qiagen). Reverse transcription was performed on 1 μg total RNA using SuperScript III (Life Technologies). To quantify allelic skewing, DNA or cDNA was amplified using the following biotinylated primers and subsequently sequenced using Q24 Pyromark (Qiagen).

### Western-blot from nuclear extract

Nuclear extracts were prepared by harvesting cells with trypsin, washing the pellet in PBS and resuspending the cells in ice-cold 10 ml buffer A (10 mM HEPES pH 7.9, 10 mM KCl, 1.5 mM MgCl2, 0.1% NP-40, c0mplete Mini Protease inhibitor EDTA free from Roche) and rotating for 10 min at 4°C. Nuclei were centrifuged at 800g for 10 min at 4 °C and resuspended in appropriate amount of RIPA buffer (50 mM Tris-HCl pH 8.0–8.5, 150 mM NaCl, 1% Triton X-100, 0.5% sodium deoxycholate, 0.1% SDS) containing c0mplete Mini Protease inhibitor (Roche), incubated for 20 min on ice and sonicated with a Bioruptor (four 5-s pulses). Lysates were then centrifuged for 30 min at 4 °C, and supernatants were kept. Protein concentration was determined using the Bradford (BioRad) assay. Samples were then boiled at 95 °C for 10 min in 3.2x LDS buffer (Thermo) containing 200 mM DTT. For RPB1, protein extracts were loaded on a 3-8% gradient gel in Tris-Acetate buffer. For RPB3, a 4-12% gel in MOPS buffer was used; as RPB3 and lamin B have very similar size and cannot be revealed on the same membrane, extracts were loaded twice on the same gel. Transfer was performed on a 0.45-μm nitrocellulose membrane using a wet-transfer system, at 350mA for 90 min at 4 °C. RPB1 membrane was cut in two so RPB1 and Lamin B (which came from the same loaded wells) were labelled separately; for RPB3 the membrane was cut in two to label one set of loaded wells for RPB3 and the other set of wells for Lamin B.

Rpb3 1:1000 with 2nd AB dilution 1:10000
Lamin B1 1:3000 with 2nd AB dilution 1:10000
Rpb1 1:500 with 2nd AB dilution 1:5000

### RNA-FISH

#### Cell preparation

Cells were dissociated using Accutase(Invitrogen), washed twice in medium, and allowed to attach on poly-l-lysine (Sigma)-coated coverslips for 10 min. Cells were fixed with 3% paraformaldehyde in PBS for 10min at room temperature, washed in PBS three times, and permeabilized with ice-cold permeabilization buffer (PBS, 0.5% Triton X-100, 2 mM vanadyl–ribonucleoside complex) for 4 min on ice, washed in 70% ethanol and stored in 70% ethanol at −20 °C or directly labelled.

#### Probes labelling and precipitation

Probes were prepared from phenol-chloroform extractions of intron-spanning bacteria artificial chromosomes (BACs) (clone RP24-157H12 for Huwe1), or plasmid (p510 for *Xist*). Probes were labelled by nick translation (Abbott) using dUTP labelled with spectrum green (Abbott) for *Huwe1* and Cy5 (Merck) for *Xist*. Labelled probes were precipitated in ethanol (3uL of probes for plasmids and 5uL for BAC, 100uL EtOH 100%, 1uL of salmon sperm DNA, 0.7uL of NaOAc 3M pH 5.2, and for BAC probes adding 4uL Cot-1 repetitive DNA), washed in 70% ethanol, dried in a speedvac at room temperature, resuspended in formamide, denatured at 75 °C for 7 min, competed at 37 °C for 1h for BAC probes with Cot-1 DNA, and quenched on ice.

#### Hybridization

Samples were dehydrated in 4 baths of increasing ethanol concentration (80%, 95%, 100% twice) and air-dried quickly. Probes were mixed in equal volume of hybridization buffer (7uL probes + 7uL of buffer: 40% dextran sulfate, 2x SSC, BSA 2mg/mL, 10 mM vanadyl-ribonucleoside), spotted on cells and hybridized at 37 °C overnight. The next day, coverslips were washed three times for 7 min with 50% formamide in 2X SSC at 42 °C, and three times for 7 min with 2X SSC. DAPI (0.2 mg ml-1) was added to the last wash and coverslips were mounted with ProLong Diamond Antifade Mountant (Invitrogen).

#### Microscopy

RNA-FISH were imaged on a OLYMPUS SpinSR10 spinning disk microscope equipped with a Yokogawa CSU-W1 unit, a UPLSAPO 100XS objective (NA 1,35, silicone oil) and using the SoRa disk (without additional magnification lens). 3D images were acquired with xx between stacks. For counting, Stacks were flatten into 2D by max projection, and cells with *Xist* clouds and/or Huwe1 foci were counted manually.

### Fluorescence Correlation Spectroscopy (FCS) and calibrated imaging

#### Cell preparation

Cells were splitted at 50,000 cells per cm2 in ibidi 8-well chamber slides with glass bottom, coated with fibronectin. The chamber slides always contained one empty well for measurement on pure dye in solution, one well of “negative control” cells (no Halo tag), and two well of “free diffusing control” Halo-NLS cells. 24h after splitting, cells were induced with doxycycline (2 μg/ml). After 24h of induction cells were labelled using Halo-ligand-JFX549 (graciously provided by Luke Lavis) in media containing doxycycline, incubated for 30min, washed 3 times in PBS, incubated 3 times in fresh media (with doxycycline) for 20min with PBS washes between incubations. For RPB1- and RPB3-Halo labelling was performed using ligand at 100nM; for Halo-NLS, as the piggyBac transgene was expressed at a much higher level, one well was labelled with 5nM and one with 2nM, to allow a large range of fluorescence intensities for the calibration. “Negative control” cells were labelled with 100nM. Media (with doxycycline) was finally changed and pure AF568 dye in solution (5nM) was added in the free well of the chamber before imaging.

#### Microscopy - FCS

FCS measurement and 3D imaging were performed on a Zeiss LSM880 microscope using a C-Apochromat Zeiss UV-visible-IR 40×/1.2-NA objective and operated with ZEN Blue software, equipped with an incubator chamber controlled at 37C and 8% CO2. FCS measurements were automatized using macro FCSRunner. The power of the 561 laser was set to 0.01 for all FCS measurements. For pure AF568 dye, 2 consecutive measurements of 10s were performed for 5 points per field of view, for at least 4 fields of view per experiment. For all measurements in cells, a single measurement of 30s was performed for a single point per cell, followed by one acquisition for the whole field of view, single stack at the same z position as the FCS measurement, with the same laser power and the following parameters: 8x zoom in and 128×128 pixels per field of view (resulting in a pixel size of 0.0991362um in x/y). All control measurements (Dye, free diffusing control and negative control) were performed each day for each individual experiment. Stage leveling was done manually based on the coverslip reflection, and re-done for each individual well of the chamber slide.

#### Microscopy - 3D acquisitions

3D acquisitions were performed on the same system as the FCS, following FCS acquisitions on the same day. Images were taken with the same parameters as FCS snapshots and 0.48um between z-stacks.

#### FCS data processing

Background average signal in negative control cells was calculated using *FCSFitM*. FCS measurements were processed using *FluctuationAnalyzer*. All parameter were kept at default value except the following:

- Step *Modify and correlate*: “Base freq”: 1,000,000 (dye measurement) or 100,000 (cell)
- Step *Intensity correction*: “Base freq”: 1,000,000 (dye measurement) or 100,000 (cell) “Offset”: 0 (Dye) or the average intensity from negative cells (cell)
- Step *Fit correlations*: all fitting were performed using the model “two-component anomalous diffusion with triplet-like blinking” with weighted fit, 2 runs of optimization and initial guess. For Dye, the fitting was performed only on lag times from 0 to 10239us to avoid overfitting the flat tail of the autocorrelation function.

#### Confocal volume estimation

the confocal volume was calculated based on FCS measurements on AF568 in solution based on the following equation:

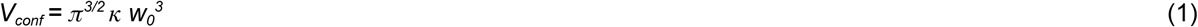

where *V_conf_* is the effective confocal volume, *k* is the ratio of axial to lateral radius of this volume (estimated from the autocorrelation fitting) and *w_0_* is the lateral radius of the confocal volume, which can be calculated following the following equation:

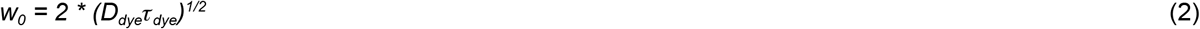

where *D_dye_* is the diffusion coefficient of the dye in solution (previously estimated to be *D_AF568_* = 521.46 um2/s at 37C (*20*)), *τ_dye_* the diffusion time of the dye (estimated from the autocorrelation fitting) and *w_0_* is the lateral radius of the confocal volume.

The average confocal volume was calculated based on all the dye measurement for each individual experiment separately.

#### Diffusion coefficient estimation

diffusion coefficients were calculated based on equation (2):

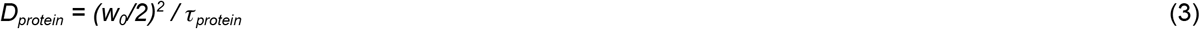

*where D_protein_* is the diffusion coefficient of the protein, *w_0_* the lateral radius of the confocal volume estimated in the previous step based on dye measurements, and *τ_protein_* the diffusion time of the protein estimated from the autocorrelation fitting. Diffusion coefficients were calculated for the first population from the fitting (i.e. the population with the highest diffusion coefficient, corresponding to the free fraction).

#### FCS-calibration

Calibration of pixel fluorescence intensity into concentration using paired 2D imaging and FCS measurements was performed using a KNIME pipeline available on gitlab: https://git.embl.de/grp-almf/FCSpipelineEMBL_KNIME Shortly, paired 2D images and FCS measurement (analyzed using *FluctuationAnalyzer* as described above) are loaded, and the fluorescence intensity in the 2D image at the coordinates of the FCS point measurement is extracted. The fluorescence background is calculated as the average of fluorescence intensities at FCS points in negative control cells (not expressing any Halo tag), and this background is subtracted from the fluorescence intensity measurements for RPB1-Halo, RPB3-Halo and Halo-NLS. A linear trend between FCS measured concentrations and background corrected intensities is then fitted using least square regression:

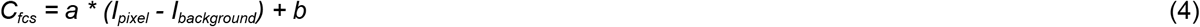

Where *C_fcs_* is the FCS measured concentration, *I_pixel_* is the fluorescence intensity at the corresponding pixel on the 2D image, and *I_background_* is the background intensity.

This calibration is then used to convert pixel fluorescence intensities in 3D images into concentration:

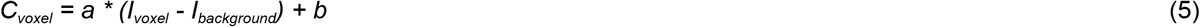

Where *C_voxel_* is the inferred concentration per voxel in 3D images, *I_voxel_* is the original fluorescence intensity per voxel for RPB1-/RPB3-Halo in 3D images, *I_background_* is the background intensity, a and b the parameters calculated in equation (4).

#### 3D images segmentation

Segmentation of nuclei, *Xist* territory, nucleoli and nucleoplasm were performed using *Ilastik* (*19*), first classifying pixels into different categories using the *autocontext* pixel classification function, and then segmenting the image based on the classifications. Different models were trained for the different regions:

- For nuclei, one model was trained using both the *Xist*-BglG-GFP and RBP1/3-Halo channels, with two annotations: background (between nuclei) and nuclei.
- For *Xist* territories, one model was trained using only the *Xist*-BglG-GFP channel; the RBP1/3-Halo was not used to not bias the segmentation of *Xist* territories based on the RPB1/3 intensities. 3 annotations were used: background, *Xist* territory and the rest of the nucleus. Only the Xist territory classification was used from this model.
- For nucleoli and nucleoplasm, a model was trained using both the *Xist*-BglG-GFP and RBP1/3-Halo channels, using 4 annotations: background, *Xist* territory (annotated as high level of *Xist*-BglG-GFP), nucleoli (annotated as low level of RBP1/3 but no *Xist*-BglG-GFP) and nucleoplasm (rest of the nucleus). The annotation of *Xist* territory was done to avoid annotating those regions as nucleoli, as they both display lower levels of RPB1/3, and are frequently spatially close; however the *Xist* territory classification from this model was not used in later analysis.

These classifications annotations were then used as input for the segmentation function (also done independently for each model).

Finally, the resulting segmentation of nuclei, *Xist* territory, nucleoli and nucleoplasm were exported in TIF format. The final segmentation was defined as follow:

- *Xist* territory: pixels belonging to ilastik segmentation of nucleus and *Xist* territory.
- nucleoli: pixels belonging to ilastik segmentation of nucleus and nucleoli but NOT *Xist* territory.
- nucleoplasm: pixels belonging to ilastik segmentation of nucleus and nucleoplasm but NOT *Xist* territory.

All ilastik models and corresponding files are available on github: https://git.embl.de/scollomb/collombet_et_al_rnapii_xist_compartment/-/tree/master/FCSCI/ilastik.

All codes for FCSCI data analysis are available on github: https://git.embl.de/scollomb/collombet_et_al_rnapii_xist_compartment/-/tree/master/FCSCI

### Single particle tracking

#### Cell labelling

Cells were splitted at 50,000 cells per cm2 in 35mm glass bottom dish (Mattek), coated with fibronectin. 24h after splitting, cells were induced with doxycycline (2 μg/ml). After 24h of induction cells were labelled using Halo-ligand-PhotoActivable-JF646 (graciously provided by Luke Lavis) at 50nM in media containing doxycycline, incubated for 30min, washed 3 times in PBS, incubated 4 times in fresh media (with doxycycline) for 30min with PBS washes between incubations. Media (with doxycycline) was finally changed before imaging.

#### Microscopy

SPT-PALM was performed as previously described REF on a custom-built Nikon TI microscope (Nikon Instruments Inc., Melville, NY) equipped with a 100x/NA 1.49 oil-immersion TIRF objective (Nikon apochromat CFI Apo TIRF 100x Oil), EM-CCD camera (Andor, Concord, MA, iXon Ultra 897; frame-transfer mode; vertical shift speed: 0.9 μs; −70°C), a perfect focusing system to correct for axial drift and motorized laser illumination (Ti-TIRF, Nikon). A 1.6x magnification lens was added in the light path allowing sampling at the objective nyquist resolution, and resulting in a pixel size of 106nm. The incubation chamber maintained a humidified 37°C atmosphere with 5% CO_2_ and the objective was also heated to 37°C. Lasers were modulated by an acousto-optic Tunable Filter (AA Opto-Electronic, France, AOTFnC-VIS-TN) and triggered with the camera TTL exposure output signal. The microscope, cameras, and hardware were controlled through NIS-Elements software (Nikon). The camera exposure time was set to 5ms, the excitation with 633nm laser (100% laser power) to 1ms and the photoactivation with 405nm laser synchronized with the off time of the camera (0.477 ms between frames). The intensity of the 405 laser was adapted manually between 2 and 10% during the acquisition to optimise photoactivation to obtain enough tracking per experiments while remaining sparse enough to track single molecules accurately. 30,000 frames were acquired per cell. Snapshots of Xist-BglG-GFP were taken before and after the SPT tracking with the 488nm laser (200ms exposure).

#### Localisation and tracking

Localisation and tracking were performed using the *pyspaz* program (https://github.com/alecheckert/pyspaz). Localisation was performed using the function *localize detect-and-localize-file* with the following parameters: *-s 1 -t 20*, and all other parameters as default. Tracking was performed using the function *track track-locs* with the following parameters: *--algorithm_type conservative --pixel_size_um 0.106 -e 3 -dm 10 -db 0.1 -f 5.477 -b 0*, and all other parameters as default.

#### Segmentation

For each acquisition, the two snapshots (before and after SPT) were combined as one multi dimensional TIF file using a custom python script. These snapshots were used for segmentation of *Xist* compartment and nucleoplasm using Ilastik. The *Auto-context* mode was first used to create probability maps, annotating pixels as *Xist* territory (high *Xist*-BglG-GFP signal), nucleoplasm (low *Xist*-BglG-GFP signal) and “background” (between nuclei). The *Tracking* mode was then used with the probability maps as input to automatically segment and annotate nuclei and *Xist* territory (one model built to track *Xist* territory, one model to track nuclei). All ilastik models and corresponding files are available on github: https://git.embl.de/scollomb/collombet_et_al_rnapii_xist_compartment/-/tree/master/SPT/ilastik.

#### Trajectories assignment to sub-compartments

assignment of trajectories to nuclei, nucleoplasm or XC was performed using a custom python script *interpolateMaskAndAssignTrajectories.py* available on our github: https://git.embl.de/scollomb/collombet_et_al_rnapii_xist_compartment/-/blob/master/SPT/interpolateMaskAndAssignTrajectories.py.

We used the following parameters: --olap_fracMin 0 --olap_fracMax 1 --pixel_subsampling_factor 1. For trajectories inside XC were defined as those for which at least one localisation was found inside the XC mask (--olap_rule any) and trajectories outside XC as those for which no localisation was found inside the mask (--olap_rule none).

#### Trajectories entering XC and control regions

control regions were created using a custom python script *interpolateMaskAndAssignTrajectories_moveMask.py* available on our github: https://git.embl.de/scollomb/collombet_et_al_rnapii_xist_compartment/-/blob/master/SPT/interpolateMaskAndAssignTrajectories_moveMask.py.

Shortly, this script takes as input the mask of nuclei and XC, randomly shift and rotate the XC mask and evaluate if the new mask respect a number of rules: the shifted mask does not overlap the original mask or a previously valid shifted mask by more than 10% of their respective size (*--maxMasksOlap 0.1*), the shifted mask is entirely inside the nucleus (*--maxMaskFracOutsideROE 0.0*) and the distance to the nuclear periphery is not different from the original mask by more than 50% (*--minMaskDistRoeDifFrac −0.5 --maxMaskDistRoeDifFrac 0.5*). If the shifted mask respects these rules it is added to the list of control regions. This operation is repeated until 10 control regions are found or 500,000 iterations are performed (we did not see the number of control regions per cell increase with higher number of iterations).

#### Bound/free fraction estimation

to estimate the bound and free fractions, a two component model was fitted to the distribution of jumps using *SpotOn* (*21*). A custom version of the python implementation of *SpotOn* was adapted to run on python 3, which can be found on our github: https://git.embl.de/scollomb/collombet_et_al_rnapii_xist_compartment/-/tree/master/SPT/Spot-On-cli.

The function *fit-and-plot-2states* was used with the following parameters: *--time_between_frames 0.5477 --gaps_allowed 0 --localisation_error 0.028 --weight_delta_t True --model_fit CDF --max_jump_length 2 --max_jumps_per_traj 3 --max_delta_t 6 --diffusion_bound_range 0,0.02 --diffusion_free_range 0.1,20*.

#### Jumps angles

The distribution of jump angles was calculated using the function *plot-jumps-angle-circular* from our implementation of SpotOn (github link) with the following parameters: *--gaps_allowed 0 --min_1dt_jump_length 0.2 --max_1dt_jump_length 3 --max_jumps_per_traj 100 --delta_t 1 --bin_width 10*.

#### Bootstrap analysis

Bootstrap was performed using the function *subsample-trajs* from our implementation of SpotOn.

All codes for SPT analysis are available on github: https://git.embl.de/scollomb/collombet_et_al_rnapii_xist_compartment/-/tree/master/SPT

### FRAP

#### Cell labelling

Cells were prepared the same way as for FCS-CI: splitted at 50,000 cells per cm2 in ibidi 8-well chamber slides with glass bottom, coated with fibronectin. 24h after splitting, cells were induced with doxycycline (2 μg/ml). After 24h of induction cells were labelled using Halo-ligand-JFX549 (graciously provided by Luke Lavis) at 100nM in media containing doxycycline, incubated for 30min, washed 3 times in PBS, incubated 3 times in fresh media (with doxycycline) for 20min with PBS washes between incubations. Media (with doxycycline) was finally changed before imaging.

#### Microscopy

FRAP was performed on the same microscope setup as FCS-CI using the same objective. Imaging was done with x18 zoom and an optimal frame size of 104 pixels, a speed of 18 corresponding to a dwelling time of 1.28us per pixel and a scan time of 31.95ms per frame. A snapshot was first taken using both 488 channel and 561 channels to visualise the *Xist* territory. Regions of interests were defined as circles of 10 pixels (to be always fully contained in the *Xist* territory) into the *Xist* territory, nucleoplasm and background (outside cells). FRAP acquisition was then performed only using the 561 channel to allow fast imaging with ~32ms between frames. Photobleaching was performed after 80 frames on the *Xist* territory circle (or a second nucleoplasm region for bleaching control in the nucleoplasm), with a scanning speed of 7 (corresponding to ~ 9ms bleaching time) and a laser poxer of 60% (both parameters were manually optimized to allow >50% bleaching inside the defined circle while minimizing bleaching of the surrounding area).

#### Data analysis

To correct for bleaching and background, an exponential decay function was fitted to the background and nucleoplasm regions measurements.

The signal in the bleached region of interest (ROI) at all time points was then scaled to the pre bleached signal, and the background was subtracted as followed:

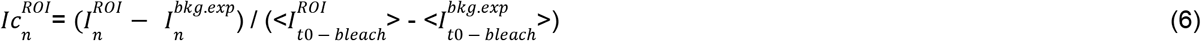

Where 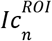 is the background and pre-bleached scaled intensity in ROI at time n, 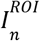 is the raw intensity in the ROI at time n, 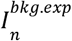 is the exponential fit of the background signal (outside cell) at time n, 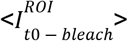 is the mean of signal in the ROI before bleaching time, and 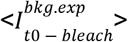 is the mean of the background fitted signal.

Bleaching was then corrected using the signal in the control region:

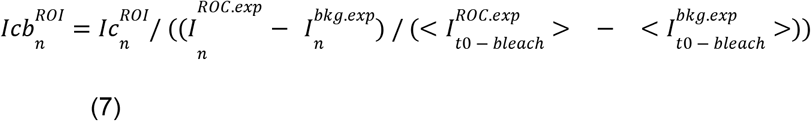

Where 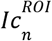 is the corrected signal in ROI at time n calculated in (6), 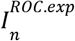 is the exponentially fitted intensity in the control region (inside the nucleus, non bleached) at time n, and 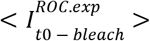 the mean of exponentially fitted intensity in the control region before bleaching.

## Supplementary figures

**Supplementary Fig S1.**
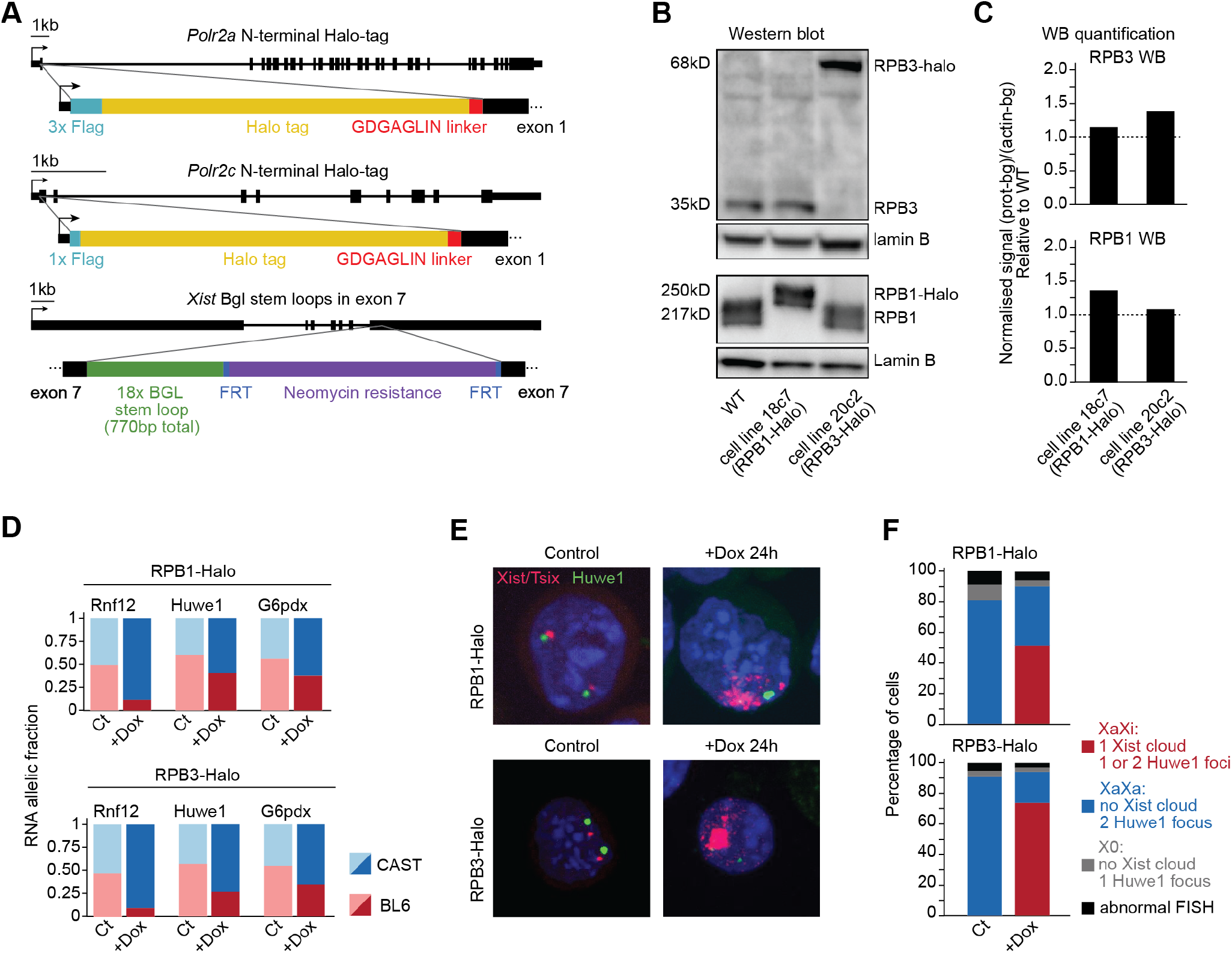
Cell line characterisation. **A.** scheme of genetic engineering for RPB1-Halo, RPB3-Halo and Xist-Bgl. **B.** Western blot for RPB1 and RPB3 in untagged cell line, RPB1-Halo and RPB3-Halo. Whole image of the WB are shown in Figure S5. **C.** Quantification of the westernblot signal from B. **D.** Allelic expression of Rnf12, Huwe1 and G6pdx measured by RNA pyrosequencing in cells before (Ct) and after 24h Xist induction. **E.** Representative examples of RNA FISH for Xist and Huwe1 in cells before (Ct) and after 24h Xist induction. **F.** Quantifications of the percentage of cells showing Xist induction (XaXi, red) no induction (XaXa, blue) or other phenotype before and after 24h Xist induction.

**Supplementary Fig S2.**
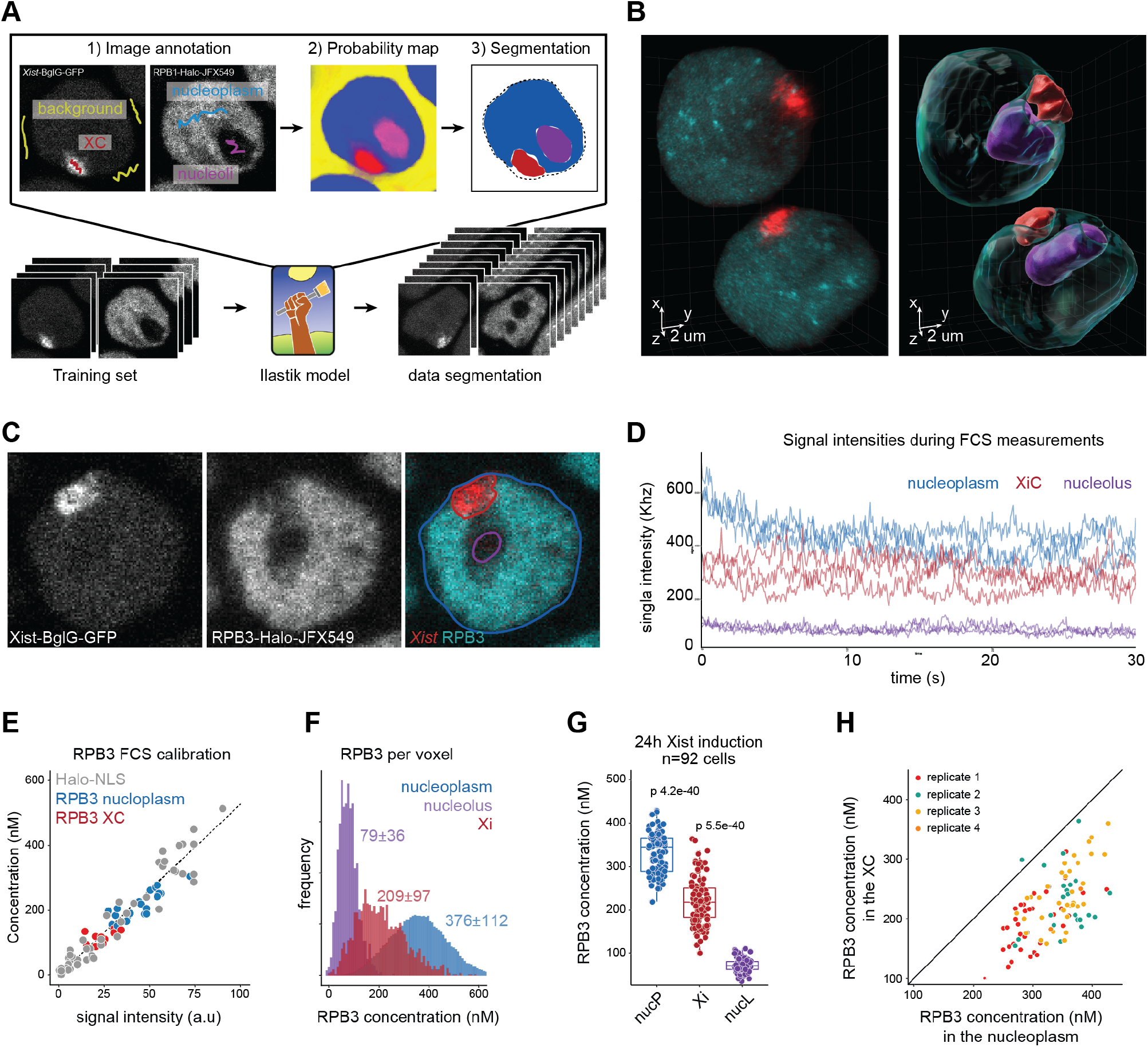
Live imaging of *Xist* RNA and RNAPII, segmentation and FCS-CI. **A.** Summary of the 3D segmentation workflow using Ilastik (*19*). **B.** 3D rendering of *Xist*-BglG-GFP and RPB1-Halo signals in live-cell confocal imaging; and of the nucleoplasm, XC and nucleolus segmentation. **C.** Representative image of confocal microscopy of *Xist*-BglG-GFP and RPB3-Halo in live cells (single Z stack) after 24h of Xist induction (doxycycline treatment) with overlaid segmentation of nucleus, nucleoli and Xi (see methods). **D.** Representative example of signal intensities and fluctuation during FCS measurement in the nucleoplasm, XC and nucleolus. **E.** Calibration of RPB3 signal intensity from point scanning imaging with FCS measured concentrations. Each dot represents a single measurement from a single cell. The linear calibration is established only on the freely diffusing Halo-NLS. **F.** Calibrated RPB3 concentration per voxel for the nucleus shown in C, based on the calibration in E. the average concentration per region is indicated (± 95% confidence interval). **G.** distribution of RPB3 average concentration per region per cell after 24h of Xist induction. Each dot represents a single cell (n=92). P-values of the differences are indicated on top (t-test two sided, paired data). **H.** RPB3 Concentration in the XC versus nucleoplasm. Each dot represents a single cell.

**Supplementary Fig S3.**
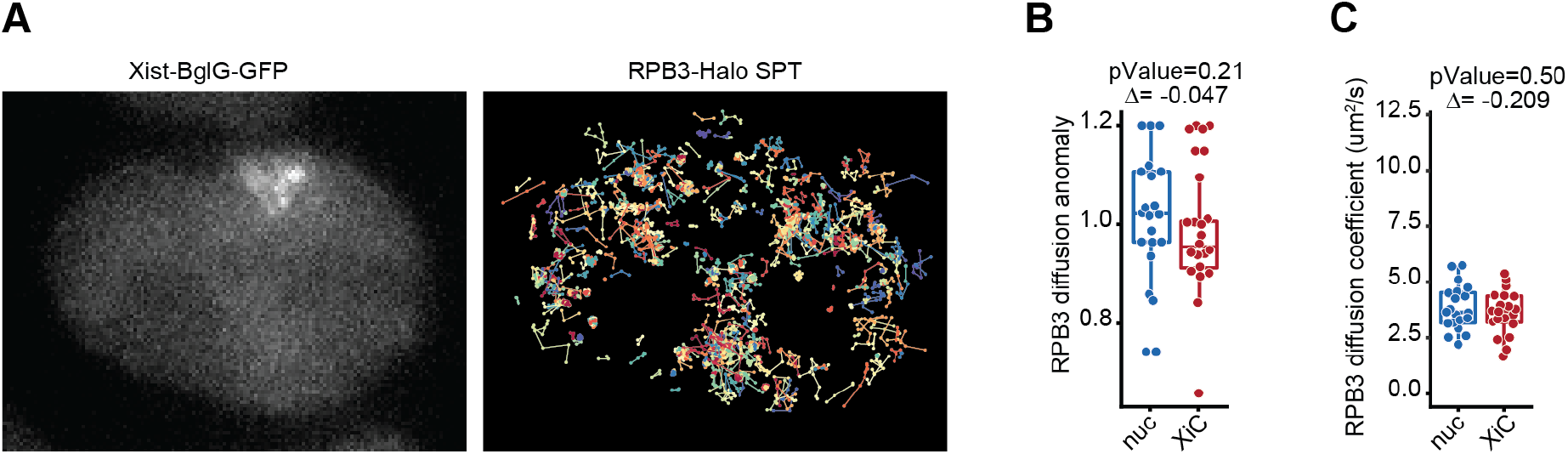
Characterisation of RNAPII diffusion on the Xi. **A.** Representative example of RPB3-Halo single particle tracking after 24h *Xist* induction. **B.** RPB3 diffusion anomaly exponent from FCS measurement inside and outside XC. Each dot represents a single cell. **C.** RPB3 diffusion coefficient inferred from FCS measurements inside and outside the XC. Each dot represents a single cell.

**Supplementary Fig S4.**
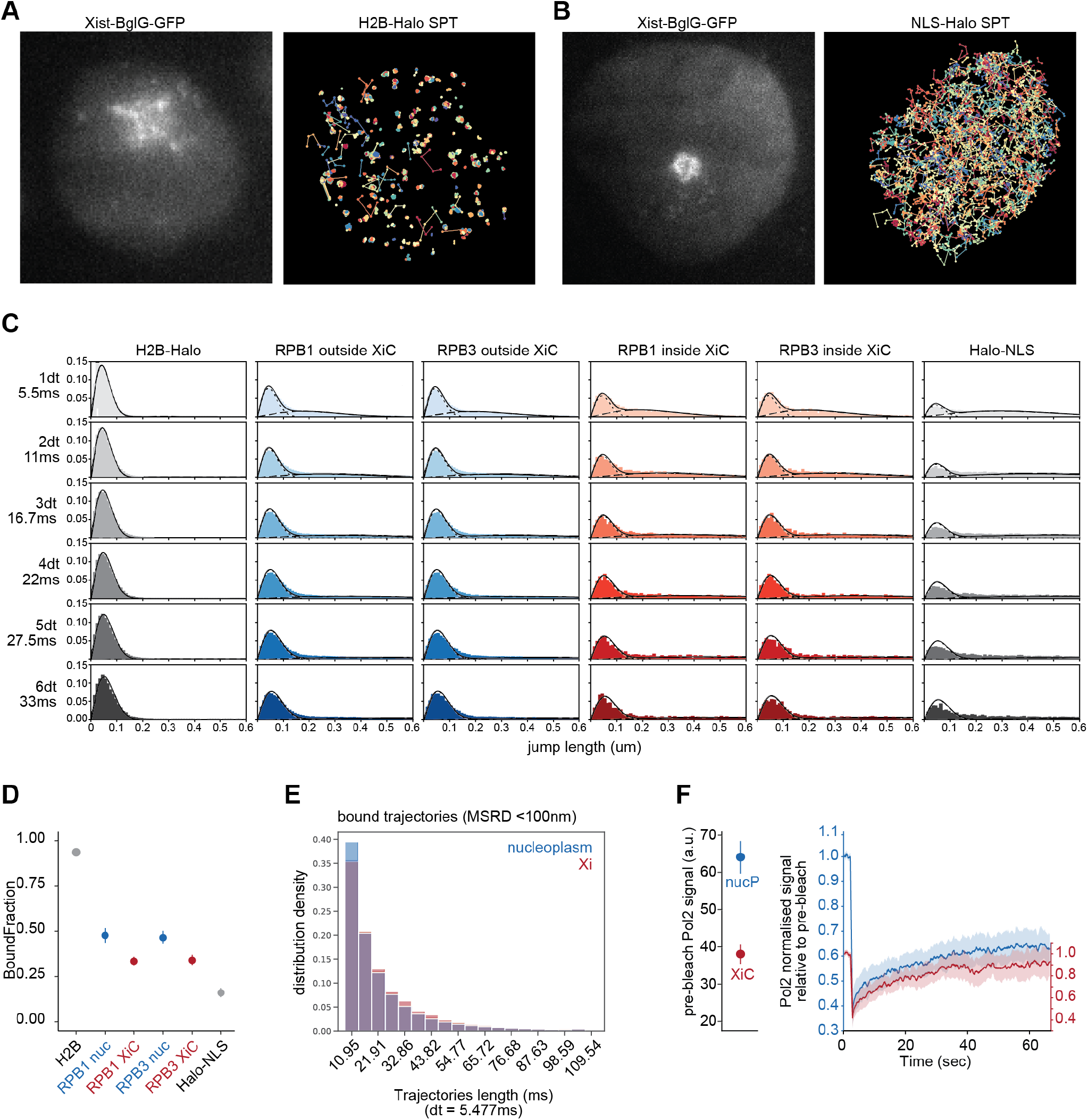
Single particle tracking and FRAP. **A.** Representative example of histone H2B-Halo single particle tracking after 24h *Xist* induction. **B.** Representative example of Nls-Halo single particle tracking after 24h *Xist* induction. **C.** Distribution of jump length in single particle tracking RPB1, RPB3 and the “bound” histone H2B-Halo and “free” Halo-NLS controls, and **D.** Estimated bound fractions. The dot represents the fraction estimated from all pooled trajectories, and the error bar the standard deviation from 50 bootstrap subsampling of 3,000 trajectories (see methods). **E.** Distribution of tracking duration for bound RPB1 molecules (MSRD<100nm) inside and outside XC. **F.** Fluorescence recovery after photobleaching (FRAP) signal as in figure 3D, but where the signal in XC is scaled to its pre-bleached intensity relative to the nucleoplasmic signal.

**Supplementary Fig S5.**
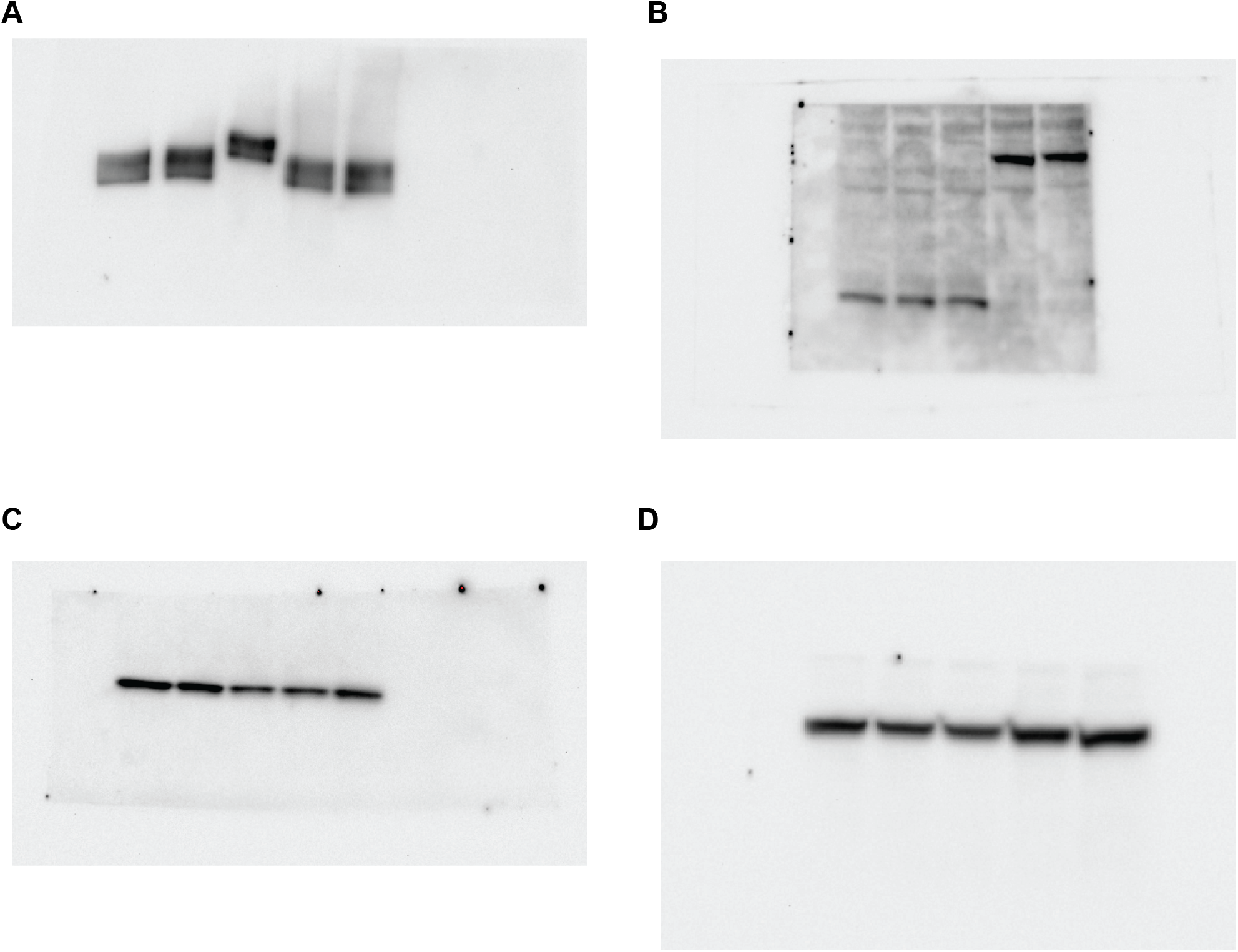
Whole image of western blots. **A.** Western Blot against RPB1. The 5 tracks are from left to right: TX1072, the original cell line in which all KI were performed; sc11c2, a cell containing only *Xist*-BglG-GFP but no RPB1/3 tagging (referred to as “WT” in Fig S1B and use as the negative control (no Halo) for FCS experiments); sc18c7, the RPB1-Halo cell line; sc20c2, the RPB3-Halo cell line used in this study; sc20c1, a second RPB3-Halo cell line not used in this study. The order of lines are the same in all western blots, and the 3 central lines (i.e. sc11c23, sc18c7 and sc20c2) are shown in Fig S1B. **B.** Western Blot against RPB3. **C**. Lamin control corresponding to the RPB1 western blot in A. The transfer membrane was cut in two to separate the migration region of RPB1 and Lamin B and labelled independently, but come but the same loaded wells (see methods). **D.** Lamin control corresponding to the RPB3 western blot in B. RPB3 and Lamin western blot were obtained from different wells of the same gel, because the two proteins have very similar sizes and their bands could not be discriminated in a single migration (see methods).

## References

1. I. Okamoto, A. P. Otte, C. D. Allis, D. Reinberg, E. Heard, Epigenetic dynamics of imprinted X inactivation during early mouse development. Science. 303, 644–649 (2004).

2. J. Chaumeil, P. Le Baccon, A. Wutz, E. Heard, A novel role for Xist RNA in the formation of a repressive nuclear compartment into which genes are recruited when silenced. Genes Dev. 20, 2223–2237 (2006).

3. P. Fraser, W. Bickmore, Nuclear organization of the genome and the potential for gene regulation. Nature. 447, 413–417 (2007).

4. J. C. Chow, C. Ciaudo, M. J. Fazzari, N. Mise, N. Servant, J. L. Glass, M. Attreed, P. Avner, A. Wutz, E. Barillot, J. M. Greally, O. Voinnet, E. Heard, LINE-1 activity in facultative heterochromatin formation during X chromosome inactivation. Cell. 141, 956–969 (2010).

5. C. A. McHugh, C.-K. Chen, A. Chow, C. F. Surka, C. Tran, P. McDonel, A. Pandya-Jones, M. Blanco, C. Burghard, A. Moradian, M. J. Sweredoski, A. A. Shishkin, J. Su, E. S. Lander, S. Hess, K. Plath, M. Guttman, The Xist lncRNA interacts directly with SHARP to silence transcription through HDAC3. Nature. 521, 232–236 (2015).

6. F. Dossin, I. Pinheiro, J. J. Żylicz, J. Roensch, S. Collombet, A. Le Saux, T. Chelmicki, M. Attia, V. Kapoor, Y. Zhan, F. Dingli, D. Loew, T. Mercher, J. Dekker, E. Heard, SPEN integrates transcriptional and epigenetic control of X-inactivation. Nature. 578, 455–460 (2020).

7. K. Plath, D. Talbot, K. M. Hamer, A. P. Otte, T. P. Yang, R. Jaenisch, B. Panning, Developmentally regulated alterations in Polycomb repressive complex 1 proteins on the inactive X chromosome. J. Cell Biol. 167, 1025–1035 (2004).

8. J. Silva, W. Mak, I. Zvetkova, R. Appanah, T. B. Nesterova, Z. Webster, A. H. F. M. Peters, T. Jenuwein, A. P. Otte, N. Brockdorff, Establishment of histone h3 methylation on the inactive X chromosome requires transient recruitment of Eed-Enx1 polycomb group complexes. Dev. Cell. 4, 481–495 (2003).

9. M. de Napoles, J. E. Mermoud, R. Wakao, Y. A. Tang, M. Endoh, R. Appanah, T. B. Nesterova, J. Silva, A. P. Otte, M. Vidal, H. Koseki, N. Brockdorff, Polycomb group proteins Ring1A/B link ubiquitylation of histone H2A to heritable gene silencing and X inactivation. Dev. Cell. 7, 663–676 (2004).

10. G. Pintacuda, G. Wei, C. Roustan, B. A. Kirmizitas, N. Solcan, A. Cerase, A. Castello, S. Mohammed, B. Moindrot, T. B. Nesterova, N. Brockdorff, hnRNPK Recruits PCGF3/5-PRC1 to the Xist RNA B-Repeat to Establish Polycomb-Mediated Chromosomal Silencing. Mol. Cell. 68, 955–969.e10 (2017).

11. H. Sunwoo, D. Colognori, J. E. Froberg, Y. Jeon, J. T. Lee, Repeat E anchors Xist RNA to the inactive X chromosomal compartment through CDKN1A-interacting protein (CIZ1). Proc. Natl. Acad. Sci. U. S. A. 114, 10654–10659 (2017).

12. A. Pandya-Jones, Y. Markaki, J. Serizay, T. Chitiashvili, W. R. Mancia Leon, A. Damianov, C. Chronis, B. Papp, C.-K. Chen, R. McKee, X.-J. Wang, A. Chau, S. Sabri, H. Leonhardt, S. Zheng, M. Guttman, D. L. Black, K. Plath, A protein assembly mediates Xist localization and gene silencing. Nature (2020), doi:10.1038/s41586-020-2703-0.

13. Y. Markaki, J. G. Chong, C. Luong, S. Y. X. Tan, Y. Wang, E. C. Jacobson, D. Maestrini, I. Dror, B. A. Mistry, J. Schöneberg, A. Banerjee, M. Guttman, T. Chou, K. Plath, Xist-seeded nucleation sites form local concentration gradients of silencing proteins to inactivate the X-chromosome, 24 (2020).

14. A. Cerase, A. Armaos, C. Neumayer, P. Avner, M. Guttman, G. G. Tartaglia, Phase separation drives X-chromosome inactivation: a hypothesis. Nat. Struct. Mol. Biol. 26, 331–334 (2019).

15. D. T. McSwiggen, M. Mir, X. Darzacq, R. Tjian, Evaluating phase separation in live cells: diagnosis, caveats, and functional consequences. Genes Dev. 33, 1619–1634 (2019).

16. E. G. Schulz, J. Meisig, T. Nakamura, I. Okamoto, A. Sieber, C. Picard, M. Borensztein, M. Saitou, N. Blüthgen, E. Heard, The two active X chromosomes in female ESCs block exit from the pluripotent state by modulating the ESC signaling network. Cell Stem Cell. 14, 203–216 (2014).

17. O. Masui, E. Heard, H. Koseki, Live Imaging of Xist RNA. Methods Mol. Biol. 1861, 67–72 (2018).

18. L. B. A. e Sousa, I. Jonkers, L. Syx, I. Dunkel, Kinetics of Xist-induced gene silencing can be predicted from combinations of epigenetic and genomic features. Genome (2019) (available at https://genome.cshlp.org/content/29/7/1087.short).

19. S. Berg, D. Kutra, T. Kroeger, C. N. Straehle, B. X. Kausler, C. Haubold, M. Schiegg, J. Ales, T. Beier, M. Rudy, K. Eren, J. I. Cervantes, B. Xu, F. Beuttenmueller, A. Wolny, C. Zhang, U. Koethe, F. A. Hamprecht, A. Kreshuk, ilastik:interactive machine learning for (bio)image analysis. Nat. Methods. 16, 1226–1232 (2019).

20. A. Z. Politi, Y. Cai, N. Walther, M. J. Hossain, B. Koch, M. Wachsmuth, J. Ellenberg, Quantitative mapping of fluorescently tagged cellular proteins using FCS-calibrated four-dimensional imaging. Nat. Protoc. 13, 1445–1464 (2018).

21. A. S. Hansen, M. Woringer, J. B. Grimm, L. D. Lavis, R. Tjian, X. Darzacq, Robust model-based analysis of single-particle tracking experiments with Spot-On. Elife. 7 (2018), doi:10.7554/eLife.33125.

22. I. Izeddin, V. Récamier, L. Bosanac, I. I. Cissé, L. Boudarene, C. Dugast-Darzacq, F. Proux, O. Bénichou, R. Voituriez, O. Bensaude, M. Dahan, X. Darzacq, Single-molecule tracking in live cells reveals distinct target-search strategies of transcription factors in the nucleus. Elife. 3 (2014), doi:10.7554/eLife.02230.

23. D. T. McSwiggen, A. S. Hansen, S. S. Teves, H. Marie-Nelly, Y. Hao, A. B. Heckert, K. K. Umemoto, C. Dugast-Darzacq, R. Tjian, X. Darzacq, Evidence for DNA-mediated nuclear compartmentalization distinct from phase separation. Elife. 8 (2019), doi:10.7554/eLife.47098.

24. R. J. Buehler, Confidence Intervals for the Product of Two Binomial Parameters. J. Am. Stat. Assoc. 52, 482–493 (1957).

25. L. Giorgetti, R. Galupa, E. P. Nora, T. Piolot, F. Lam, J. Dekker, G. Tiana, E. Heard, Predictive polymer modeling reveals coupled fluctuations in chromosome conformation and transcription. Cell. 157, 950–963 (2014).

26. S. Collombet, N. Ranisavljevic, T. Nagano, C. Varnai, T. Shisode, W. Leung, T. Piolot, R. Galupa, M. Borensztein, N. Servant, P. Fraser, A. Katia, E. Heard, Parental-to-embryo switch of chromosome organization in early embryogenesis. Nature. in press (2020).

27. D. Smeets, Y. Markaki, V. J. Schmid, F. Kraus, A. Tattermusch, A. Cerase, M. Sterr, S. Fiedler, J. Demmerle, J. Popken, H. Leonhardt, N. Brockdorff, T. Cremer, L. Schermelleh, M. Cremer, Three-dimensional super-resolution microscopy of the inactive X chromosome territory reveals a collapse of its active nuclear compartment harboring distinct Xist RNA foci. Epigenetics Chromatin. 7, 8 (2014).

28. A. Rego, P. B. Sinclair, W. Tao, I. Kireev, A. S. Belmont, The facultative heterochromatin of the inactive X chromosome has a distinctive condensed ultrastructure. J. Cell Sci. 121, 1119–1127 (2008).

29. B. G. O’Flynn, T. Mittag, The role of liquid–liquid phase separation in regulating enzyme activity. Curr. Opin. Cell Biol. 69, 70–79 (2021).

30. A. Wutz, T. P. Rasmussen, R. Jaenisch, Chromosomal silencing and localization are mediated by different domains of Xist RNA. Nat. Genet. 30, 167–174 (2002).

31. J. M. Engreitz, A. Pandya-Jones, P. McDonel, A. Shishkin, K. Sirokman, C. Surka, S. Kadri, J. Xing, A. Goren, E. S. Lander, K. Plath, M. Guttman, The Xist lncRNA exploits three-dimensional genome architecture to spread across the X-chromosome. Science. 341, 1237973 (2013).

32. X. Darzacq, Y. Shav-Tal, V. de Turris, Y. Brody, S. M. Shenoy, R. D. Phair, R. H. Singer, In vivo dynamics of RNA polymerase II transcription. Nat. Struct. Mol. Biol. 14, 796–806 (2007).

33. K. Teller, D. Illner, S. Thamm, C. S. Casas-Delucchi, R. Versteeg, M. Indemans, T. Cremer, M. Cremer, A top-down analysis of Xa- and Xi-territories reveals differences of higher order structure at ≥ 20 Mb genomic length scales. Nucleus. 2, 465–477 (2011).

34. M. R. Gdula, T. B. Nesterova, G. Pintacuda, J. Godwin, Y. Zhan, H. Ozadam, M. McClellan, D. Moralli, F. Krueger, C. M. Green, W. Reik, S. Kriaucionis, E. Heard, J. Dekker, N. Brockdorff, The non-canonical SMC protein SmcHD1 antagonises TAD formation and compartmentalisation on the inactive X chromosome. Nat. Commun. 10, 30 (2019).

35. C. Y. Wang, T. Jégu, H. P. Chu, H. J. Oh, J. T. Lee, SMCHD1 Merges Chromosome Compartments and Assists Formation of Super-Structures on the Inactive X. Cell, 1–16 (2018).

